# Group size and collective action in a binary contribution game

**DOI:** 10.1101/483149

**Authors:** Georg Nöldeke, Jorge Peña

## Abstract

We consider how group size affects the private provision of a public good with non-refundable binary contributions. A fixed amount of the good is provided if and only if the number of contributors reaches an exogenous threshold. The threshold, the group size, and the identical, non-refundable cost of contributing to the public good are common knowledge. Our focus is on the case in which the threshold is larger than one, so that teamwork is required to produce the public good. We show that both expected payoffs and the probability that the public good is obtained in the best symmetric equilibrium are decreasing in group size. We also characterize the limit outcome when group size converges to infinity and provide precise conditions under which the expected number of contributors is decreasing or increasing in group size for sufficiently large groups.

**JEL classification:** C72, D71, H41.

## 1. Introduction

Ever since the publication of Olson’s “The Logic of Collective Action” (Olson, 1965), group size has been considered important in determining how successful a group will be in attaining its common goals. Specifically, Olson suggested that “[t]he larger a group is, the farther it will fall short of providing the optimal supply of any collective good, and the less likely that it will act to obtain even a minimal amount of such a good. In short, the larger the group, the less it will further its common interests” (Olson, 1965, p. 36). These suggestions have attracted much attention in economics and political science but providing firm theoretical underpinnings has proven challenging (Sandler, 2015). While such group size effects are well understood for some of the standard models of collective action (e.g., Chamberlin, 1974; McGuire, 1974; Andreoni, 1988), for other such models this is not the case.

In this paper we investigate group size effects for the class of participation games without refunds introduced in Palfrey and Rosenthal (1984) to model the private provision of a discrete public good. In such a threshold game, *n* group members decide simultaneously whether to contribute to a public good or not (to participate or not). All contributors pay a non-refundable cost. The public good is provided if and only if the number of contributors reaches an exogenous threshold *k*, in which case all group members receive the same benefit from the provision of the public good. The threshold, the group size, and the identical cost of contributing to the public good are known to all players. We assume that the threshold is larger than one, thereby focusing on what Myatt and Wallace (2008b) call a “teamwork dilemma” rather than on the volunteer’s dilemma popularized by Diekmann (1985).

This threshold game is a stark model, which deliberately abstracts from asymmetries and incomplete information about costs and benefits by assuming that all group members are identical (Palfrey and Rosenthal, 1984, p. 172). For our purposes this is an attractive features as it isolates the effects of changes in group size from the effects of changes in the composition of the group, thereby allowing us to focus on the former.^1^ We further abstract from the possibility of role identification and thus restrict attention to symmetric (Nash) equilibria in which all group members employ the same (mixed) strategy and therefore obtain the same equilibrium payoff.^2^ Of course, the assumption that there is no role identification makes what is already a stark model even starker. In particular, this assumption is not satisfied if group members interact repeatedly and therefore can condition their behavior on actions played in the past. Our approach to the investigation of group size effects is thus orthogonal to the one pursued in Myatt and Wallace (2002), Myatt and Wallace (2008a) and Myatt and Wallace (2008b) who use stochastic stability arguments akin to the ones developed in Kandori et al. (1993) and Young (1993) to select among the pure strategy equilibria in repeated threshold games played by myopic players and study the composition affects arising in asymmetric games together with group size effects. Due to their focus on pure strategy equilibria, the papers by Myatt and Wallace find that the selected equilibrium features a probability of provision that is either zero or one. In contrast, we model a situation in which the probability of provision can change more gradually with group size. Over a large range of parameter values this is indeed what we find.

Generically (more precisely, for all but a countable set of cost parameters) our model has one or three symmetric equilibria. As it is always a symmetric equilibrium for no one to contribute, uniqueness obtains if and only if cost is so high and the group so large as to imply that the public good cannot be provided in a symmetric equilibrium. Hence, we face a multiplicity problem whenever the model allows for a non-trivial symmetric equilibrium, that is, an equilibrium in which the public good is provided with strictly positive probability. We resolve this problem by focusing on the — uniquely determined — best symmetric equilibria, that is, the one which provides all agents with the highest expected payoff among the symmetric equilibria. This is also the symmetric equilibrium with the highest participation probability and the highest success probability (i.e., the highest probability that the public good is provided) among the symmetric equilibria and we thus refer to it as the *maximal equilibrium*.

We can think of two distinct reasons justifying our focus on the maximal equilibrium. First, it is of intrinsic economic interest to ask how the best equilibrium outcome that a group can achieve (under the coordination friction implied by the symmetry constraint that we presume to be unavoidable) depends on group size. Second, among the two non-trivial symmetric equilibria the one we consider is stable under the standard dynamics (e.g., the replicator dynamics) considered in evolutionary game theory, whereas the other one is unstable.^3^ Consequently, whenever there are multiple symmetric equilibria only the maximal and the trivial equilibrium (which is always stable) are of relevance. In an evolutionary analysis of threshold games it is then natural to focus on the comparative statics of the maximal equilibrium (cf. Bach et al., 2006; Archetti and Scheuring, 2011). Nevertheless, it is of interest to understand how our comparative statics results depend on our focus on the maximal equilibrium. We therefore discuss how some of our key results would change if one were to consider the smaller of the two non-trivial symmetric equilibria instead.

Section 2 notes some fundamental facts about the symmetric equilibria of the threshold game we consider. All of these are immediate from Palfrey and Rosenthal (1984). We then begin our study of maximal equilibria by observing that (over the range of group sizes for which a non-trivial symmetric equilibrium exists) the participation probability in such an equilibrium is strictly decreasing in group size. We state this well-known result (e.g., Offerman, 1997; Hindriks and Pancs, 2002) as Proposition 1 in Section 3.1.

Our main results (Proposition 2 in Section 3.2 and Proposition 3 in Section 3.3) show that the expected payoff and success probability that are induced by the maximal equilibrium are strictly decreasing in group size until group size reaches a critical value (which may be infinite) beyond which the public good cannot be provided. We view Proposition 2 as confirmation of Olson’s maxim that “the larger the group, the less it will further its common interests,” whereas Proposition 3 formalizes his statement that the larger a group is, “the less likely that it will act to obtain even a minimal amount of such a good.” We complement these results by characterizing the limit as group size goes to infinity (Proposition 4 in Section 3.4) and by showing that for sufficiently large groups the expected number of participants is increasing in group size if and only if the contribution cost is sufficiently low (Proposition 5 in Section 3.5). We find the latter result interesting as it delineates the circumstances under which larger groups not only have a lower success probability but also incur higher expected aggregate costs than smaller groups.

The questions addressed in our Propositions 3 and 5 have been previously considered in Hindriks and Pancs (2002).^4^ In their Proposition 6 these authors claim that “the effect of group size on the probability of provision is indeterminate.” In contrast, our Proposition 3 shows that this effect can be determined and is negative for the maximal equilibrium we consider. Hindriks and Pancs (2002, Proposition 7) also consider the case of large group sizes. While three out of the four claims in that proposition are (as we prove) correct, their argument yielding these results is not. The problem is that Hindriks and Pancs (2002) take for granted that the error introduced by using the Poisson approximation to the binomial distribution can be ignored when studying comparative statics for sufficiently large groups. Our result on the expected number of contributors — which directly contradicts the corresponding claim in Proposition 7 from Hindriks and Pancs (2002) — shows that this is not so.

As we have noted before, the games we consider differ from the volunteer’s dilemma only in that we consider thresholds *k* > 1, whereas the volunteer’s dilemma corresponds to *k* = 1. The kind of group size effects we study are much easier to determine for the volunteer’s dilemma (see, for instance, the textbook treatment in Dixit et al. 2004, p. 454–458) because its unique symmetric equilibrium is easy to calculate explicitly. With the exception of Proposition 2 — in the volunteer’s dilemma the payoff in the symmetric equilibrium is independent of group size — all our propositions generalize corresponding results for the symmetric equilibrium of the volunteer’s dilemma.

We discuss some other related literature that considers symmetric equilibria in symmetric participation games to study group size effects in Section 4, where we also note how our Propositions 1 – 3 can be extended to the second class of games considered in Palfrey and Rosenthal (1984), namely participation games with refunds.

## 2. The threshold game

We consider the complete-information participation game from Palfrey and Rosenthal (1984) with non-refundable contributions. For simplicity, we refer to this as the threshold game. We first present the model and then collect some facts that are relevant for the study of its symmetric equilibria. These facts are immediate from the analysis in Palfrey and Rosenthal (1984), but for our subsequent analysis it will be convenient to state them here.

### 2.1. Model

There is a group of *n* > 2 players. Players simultaneously decide whether or not to participate in the provision of a public good by making a fixed contribution. If *k* or more group members participate, the public good is provided. Otherwise, the public good is not provided. Every player derives a benefit, which we normalize to one, if the public good is provided and pays the participation cost *c* if participating. Payoffs are the difference between the benefit obtained and the cost incurred. The participation cost satisfies 0 < *c* < 1.

For *k* = 1 this game is the *volunteer’s dilemma* popularized by Diekmann (1985) in which one contribution suffices to ensure the provision of the public good. As the volunteer’s dilemma is well understood, we assume 1 < *k* < *n* throughout the following.^5^

### 2.2. Symmetric strategy profiles and equilibria

As explained in the Introduction, we focus on the symmetric Nash equilibria of the threshold game.^6^

We identify any symmetric strategy profile with the common *participation probability p* ∈ [0, 1] and let

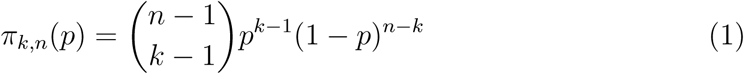

denote the probability that any player is pivotal for the provision of the public good when such a strategy profile is played: as the participation of a player will make the difference between provision and non-provision of the public good if and only if *k* − 1 out of the *n* − 1 other players participate, this probability is given by the binomial expression on the right side of (1).

As we have assumed 1 < *k* < *n*, it is clear that *p* = 0 is a symmetric equilibrium whereas *p* = 1 is not. Further, a symmetric strategy profile with 0 < *p* < 1 is a symmetric equilibrium if and only if players are indifferent between participating or not. This indifference condition requires the probability *π*_*k,n*_(*p*) that a player is pivotal to be equal to the participation cost:

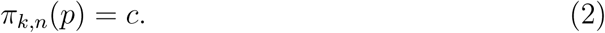

It is easily verified that the pivot probability *π*_*k,n*_(*p*) has the following *unimodality properties*: (i) it is differentiable in *p*, (ii) it satisfies *π*_*k,n*_(0) = *π*_*k,n*_(1) = 0, and (iii) it is strictly increasing on the interval [0, (*k* − 1)/(*n* − 1)] and strictly decreasing on the interval [(*k* − 1)/(*n* − 1), 1] with non-zero derivative on the interiors of these intervals.^7^ In particular, (*k* − 1)/(*n* − 1) is the unique maximizer of the pivot probability *π*_*k,n*_(*p*) in the interval [0, 1]. Hence,

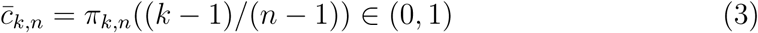

is the critical value of the participation cost such that for costs above this level the pivotality condition (2) has no solution and, thus, no interior symmetric equilibrium exists. If 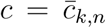 holds, then (*k* − 1)/(*n* − 1) is the unique solution to (2). If 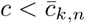 holds, the unimodality properties of *π*_*k,n*_(*p*) imply that (2) has one solution to the left and one solution to the right of (*k* − 1)/(*n* − 1). This gives the following characterization result for the number and location of symmetric strategy equilibria, which specializes the characterization results in Palfrey and Rosenthal (1984, Section to the symmetric equilibria under consideration here. Figure 1 illustrates.

**Figure 1:**
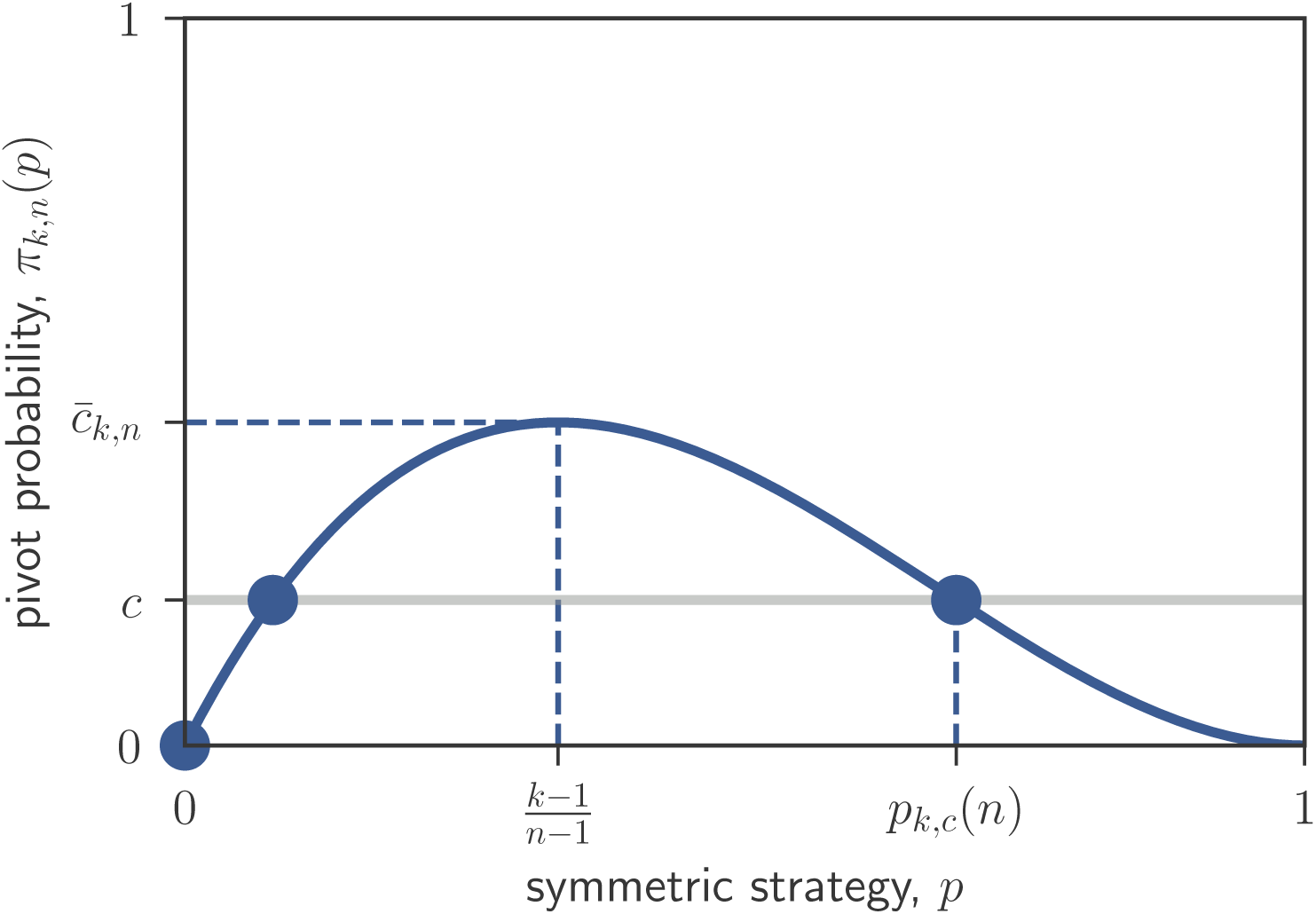
Symmetric equilibria (*circles*) and the pivot probability (*solid curve*), here illustrated for *k* = 2, *n* = 4, and *c* = 0.2. The pivot probability *π*_*k,c*_(*p*) is unimodal and has a unique maximum at (*k* − 1)/(*n* − 1) with corresponding pivot probability 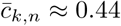. For 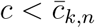 there are three symmetric equilibria with the maximal equilibrium (see Section 2.3), denoted by *p*_*k,c*_(*n*), satisfying *p*_*k,c*_(*n*) *>* (*k* − 1)/(*n* − 1).

#### Lemma 1.

*For* 1 < *k* < *n the number and location of symmetric equilibria depends on c as follows:*

1. *If 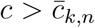, then p* = 0 *is the unique symmetric equilibrium.*
2. *If 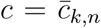, then there are two symmetric equilibria, namely p* = 0 *and p* = (*k* − 1)/(*n* − 1).
3. *If 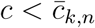, then there are three symmetric equilibria, namely p* = 0, *the unique solution to* (2) *in the interval* (0, (*k* − 1)/(*n* − 1)), *and the unique solution to* (2) *in the interval* ((*k* − 1)/(*n* − 1), 1).

We will refer to the symmetric equilibrium with *p* = 0 as the *trivial equilibrium* and to symmetric equilibria with *p* > 0 as *non-trivial equilibria*. The following result shows how the critical cost level 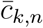 from equation (3), which determines whether non-trivial equilibria exist, depends on group size. The straightforward proof is in Appendix A.1.

#### Lemma 2.

*For any k* > 1, *the sequence 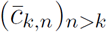 is strictly decreasing with limit*

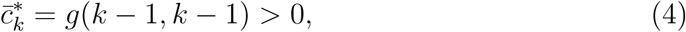

*where*

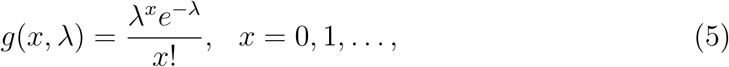

*denotes the probability mass function of a Poisson distribution with parameter λ* > 0.

To simplify the exposition, we exclude the uninteresting case in which the trivial equilibrium is the unique equilibrium for all group sizes and the non-generic case in which there exists a unique non-trivial equilibrium for some group size. That is, throughout the following we impose

#### Assumption 1.

*For any k* > 1, *the cost parameter c satisfies*

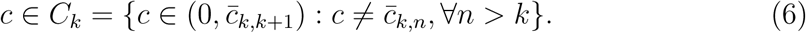

It is clear from Lemmas 1 and 2 that under Assumption 1 there exists 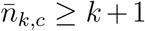 such that two non-trivial equilibria exist if and only if 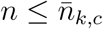 (where 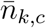 is infinite if and only if 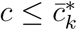) and the trivial equilibrium is the unique symmetric equilibrium otherwise. We refer to 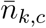 as the *critical group size*.

#### Remark 1.

Throughout our analysis we consider changes in group size for fixed threshold *k*. A natural alternative, briefly considered in Palfrey and Rosenthal (1984), is to suppose that the threshold is proportional to group size. For this case Palfrey and Rosenthal (1984, p. 178) argue that “the only equilibria which are supported by positive contribution costs for all *n* are the pure strategy equilibria”, that is, the counterpart to our critical group size 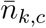 is always finite. More generally, Palfrey and Rosenthal (1984) suggest (on the same page) that “mixed strategy equilibria seem to ‘disappear’ in large populations.” In our model this is captured by the observation (see Proposition 4) that the participation probability in the sequence of maximal symmetric equilibria (and thus, a fortiori, for all sequences of symmetric equilibria) converges to zero as group size diverges to infinity. As Proposition 4 also shows, this does not imply that the equilibrium payoff and the success probability converge to zero.

### 2.3. Success probability, expected payoff, and the maximal equilibrium

Let

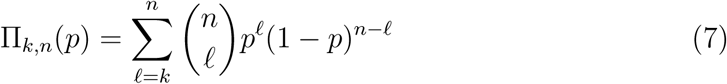

be the *success probability*, that is, the probability that the public good is provided in the symmetric strategy profile *p*. If *p* is a symmetric equilibrium, then it is a best response for any group member to choose not to participate, so that the expected payoff of every group member satisfies

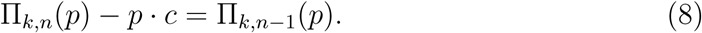

As Π_*k,n*−1_(*p*) and Π_*k,n*_(*p*) are both strictly increasing in *p* (Lehmann and Romano, 2005, Chapter 3.4) it follows that among the symmetric equilibria for given parameter values (*k, c, n*), the one with the highest participation probability *p* also induces the highest equilibrium payoff and the highest success probability. We thus refer to this equilibrium as the *maximal equilibrium*. We denote it by *p*_*k,c*_(*n*) and let

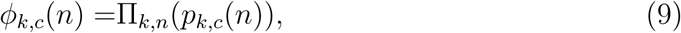

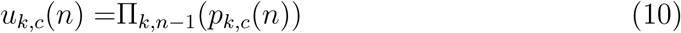

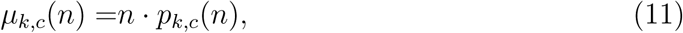

denote the success probability, expected payoff, and expected number of contributors at this maximal equilibrium.

By definition of the critical group size 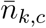 we have

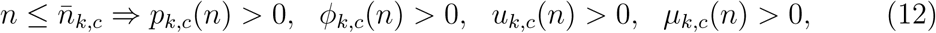

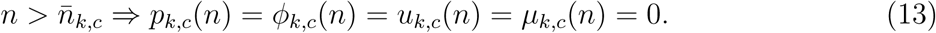

Our interest in the following is to characterize, for given (*k, c*), the comparative statics of *p*_*k,c*_(*n*), *ϕ*_*k,c*_(*n*), and *u*_*k,c*_(*n*) over the range of group sizes in (12). For the case 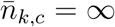 we also investigate the limit as group size converges to infinity and characterize the monotonicity properties of *µ*_*k,c*_(*n*) for large groups.

## 3. Results

In Sections 3.1 – 3.3 we establish that over the relevant range of group sizes the participation probability *p*_*k,c*_(*n*), the expected payoff *u*_*k,c*_(*n*), and the success probability *ϕ*_*k,c*_(*n*) are all strictly decreasing in group size *n*. Figure 2 illustrates (for an example with finite 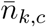). By far the deepest of these results is Proposition 2 in Section 3.2, which establishes the monotonicity of the expected payoff. The result that the success probability *ϕ*_*k,c*_(*n*) is strictly decreasing in *n* (Proposition 3 in Section 3.1) is obtained as an immediate consequence of Proposition 2 and the familiar observation, here restated as Proposition 1 in Section 3.1, that the participation probability is strictly decreasing in *n*.

**Figure 2:**
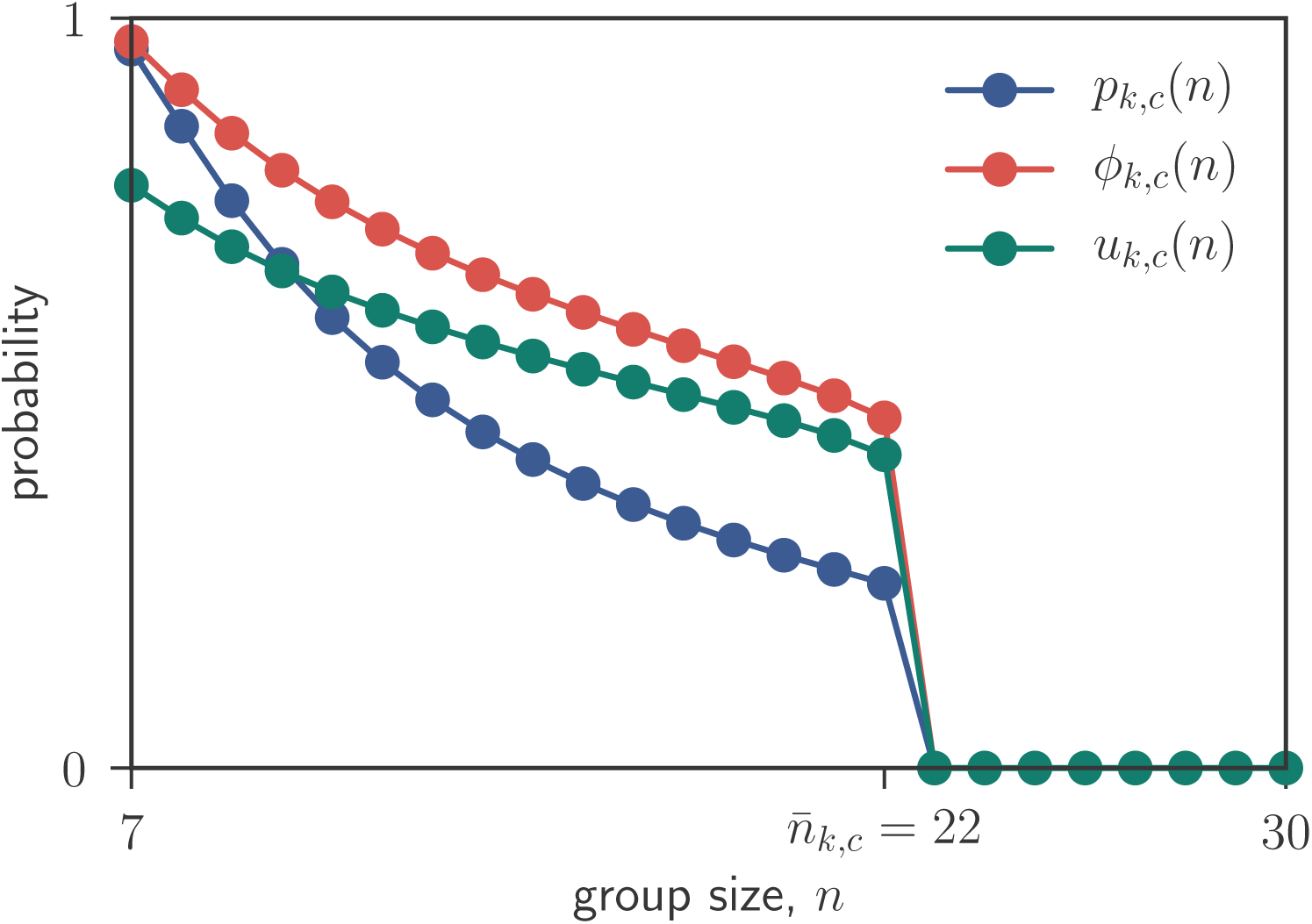
Illustration of Propositions 1 – 3 for *k* = 6 and *c* = 0.2. The participation probability *p*_*k,c*_(*n*), the equilibrium payoff *u*_*k,c*_(*n*), and the success probability *ϕ*_*k,c*_(*n*) are all strictly positive and strictly decreasing up to the critical group size 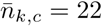. For larger groups *p*_*k,c*_(*n*), *u*_*k,c*_(*n*), and *ϕ*_*k,c*_(*n*) are all zero.

We record limit results for *n* → ∞ in Section 3.4, showing that the success probabilities, expected payoffs and expected number of contributors induced by a sequence of maximal equilibria converge to their counterparts in a game in which group sizes is Poisson distributed as in Makris (2009). Finally, building on these limit results, Section 3.5 identifies the precise conditions under which the expected number of contributors is decreasing, resp. increasing in group size for sufficiently large *n*.

### 3.1. The effect of group size on the participation probability

The maximal participation probability that can be sustained in a symmetric equilibrium is strictly decreasing in group size for 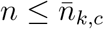 (with (13) ensuring that it drops to zero thereafter):

#### Proposition 1.

*Let 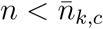. Then p*_*k,c*_(*n* + 1) < *p*_*k,c*_(*n*) *holds.*

This result has been noted before by Offerman (1997, Theorem 2.3) and Hindriks and Pancs (2002, Proposition 6(i)). We provide a proof in Appendix A.2 to make the paper self-contained and to prepare the ground for the proof of Proposition 2. Figure 3 illustrates graphically how Proposition 1 results from the relationship between the pivot probabilities *π*_*k,n*_(*p*) and *π*_*k,n*+1_(*p*). Before proceeding, we note that, as suggested by Figure 3, Proposition 1 would also hold if we were to consider the comparative statics of the smaller of the two non-trivial equilibria.^8^

**Figure 3:**
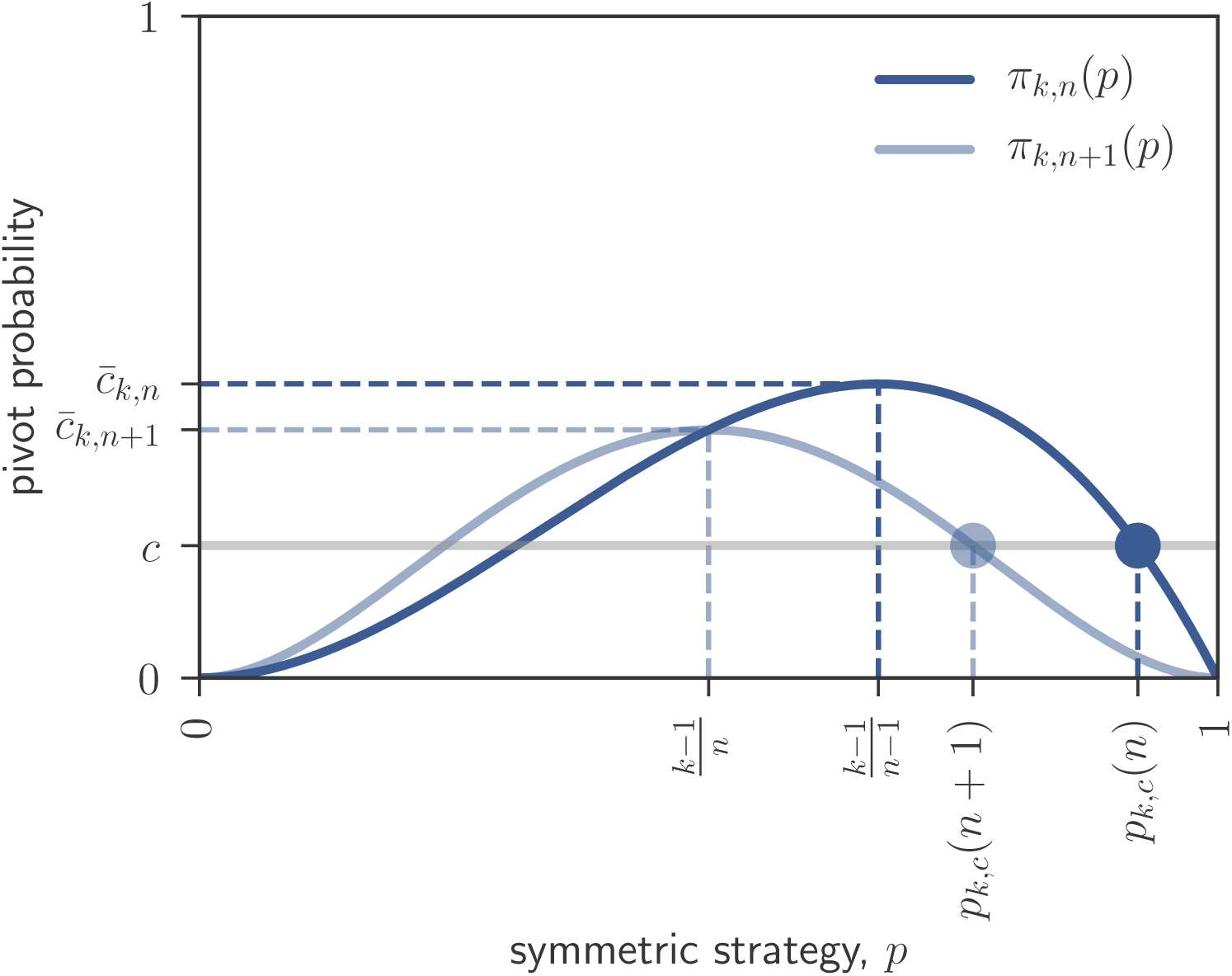
Illustration of Proposition 1 for *k* = 3, *n* = 4, and *c* = 0.2. The pivot probability *π*_*k,n*+1_(*p*) lies below *π*_*k,n*_(*p*) to the right of (*k* − 1)*/n*, implying that the maximal equilibria for consecutive group sizes satisfy *p*_*k,c*_(*n* + 1) < *p*_*k,c*_(*n*).

### 3.2. The effect of group size on the equilibrium payoff

As we have noted in Section 2.3, in any symmetric equilibrium the payoff of every group member is the same as if they were to choose non-participation while the other group members follow the equilibrium strategy. Hence, the payoff in the maximal equilibrium is, as stated in equation (10), given by *u*_*k,c*_(*n*) = Π_*k,n*−1_(*p*_*k,c*_(*n*)). It is then clear from Proposition 1 that there are two countervailing effects of an increase in group size on *u*_*k,c*_(*n*): First, increasing group size for a given participation probability *p* raises the probability that at least *k* of the *n* − 1 other group members provide and therefore has a positive effect on the equilibrium payoff. Second, we know from Proposition 1 that an increase in group size does not leave the participation probability unchanged but decreases it. For given group size this causes the probability that at least *k* of the other group members contribute to fall, and therefore has a negative effect on the equilibrium payoff. The following result shows that the latter effect dominates:

#### Proposition 2.

*Let 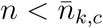. Then u*_*k,c*_(*n* + 1) < *u*_*k,c*_(*n*) *holds.*

The proof of Proposition 2 is in Appendix A.3. It proceeds in two steps. The first step shows that the equality *π*_*k,n*+1_(*p*_*k,c*_(*n* + 1)) = *π*_*k,n*_(*p*_*k,c*_(*n*)), which holds by the pivotality condition (2), implies that for the maximal equilibrium the probability that exactly *k* out of *n* − 1 group members participate is decreasing in group size, that is,

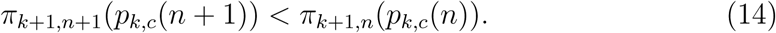

The second step infers from inequality (14) that the probability that at most *k* − 1 out of *n* − 1 group members participate is strictly increasing in group size, so that the complementary probability that at least *k* group members participate, Π_*k,n*−1_(*p*_*k,c*_(*n*)), is strictly decreasing in group size. Both steps of the proof rely on a unimodality property of the ratio of binomial probabilities with two different sample sizes from Klenke and Mattner (2010).

It is essential for the result in Proposition 2 that we consider the maximal equilibria for group sizes *n* and *n* + 1. Indeed, arguments entirely analogous to the ones proving Proposition 2 show that considering the smaller of the two solutions to the pivotality condition for both group sizes reverses the inequality in (14) and thereby also reverses the result: the payoff in these equilibria is strictly *increasing* in group size.

### 3.3. The effect of group size on the success probability

Using (8) and (9), we can rewrite (10) as

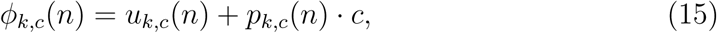

thereby decomposing the effect of group size on the success probability induced by the maximal equilibrium into two effects, namely the effect on the equilibrium payoff and the effect on the participation probability. From Propositions 1 and 2 both of these effects point in the same direction. Hence, the following proposition, is now immediate:

#### Proposition 3.

*Let 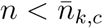. Then ϕ*_*k,c*_(*n* + 1) < *ϕ*_*k,c*_(*n*) *holds.*

The result in Proposition 3 is in sharp contrast to the corresponding claim in Proposition 6 of Hindriks and Pancs (2002), who assert that “the effect of group size on the probability of provision is indeterminate.” There are two sources for this divergence in results. First, Hindriks and Pancs (2002) actually do not show that the effect is indeterminate but simply observe that there are two countervailing effects of an increase in group size on the success probability (probability of provision). In contrast, our Proposition 3 establishes that for the maximal equilibrium the negative effect of an increase in group size on the participation probability dominates the positive effect of having more potential contributors. Second, we restrict attention to maximal equilibria, whereas Hindriks and Pancs (2002) do not impose an explicit equilibrium selection rule. This matters because, as we have noted in the last paragraphs of Sections 3.1 and 3.2, the result from Proposition 1 is unchanged but the result from Proposition 2 is reversed when considering the smaller of the two non-trivial equilibria, so that the argument proving Proposition 3 is no longer applicable. Indeed, as we have verified numerically, depending on parameter values the success probability induced by the smaller of the two non-trivial equilibria can either decrease or increase when group size is changed, so that for this equilibrium the claim in Proposition 6 of Hindriks and Pancs (2002) is correct.

### 3.4. The Poisson limit

From the monotonicity results in Sections 3.1 – 3.3 the limits

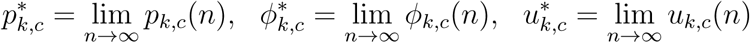

are all well defined. As we will see below, the same is true for the limit of the expected number of contributors,

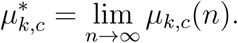

In the following results, we consider the case in which the maximal equilibrium is non-trivial for all group sizes, that is, we suppose 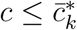 (see Lemma 2).^9^

Define

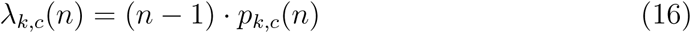

for all *n* > *k*. From the perspective of each player, *λ*_*k,c*_(*n*) is the expected number of other group members that will contribute in the maximal equilibrium *p*_*k,c*_(*n*). Recall that we have used *g*(*x, λ*) to denote the probability mass function of a Poisson distribution with expected value *λ* (see equation (5)). Appendix A.4 proves:

#### Lemma 3.

*Let 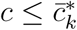. Then 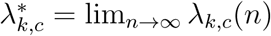 is given by the unique solution to the condition g*(*k* − 1, *λ*) = *c that satisfies λ* ≥ *k* − 1.

The condition *g*(*k* − 1, *λ*) = *c* in the statement of Lemma 3 is the natural counterpart to the pivotality condition (2) when the number of other contributors follows a Poisson distribution with expected value *λ*. For 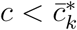, this condition has two solutions. Lemma 3 indicates that the limit value 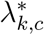 of the expected number of other contributors is given by the larger of those two solutions. This is analogous to Lemma 1 identifying non-trivial maximal equilibria as the larger of the two solutions to the pivotality condition (2). This solution satisfies 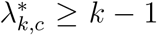 because (from Lemma 1) the inequality *λ*_*k,c*_(*n*) > *k* − 1 holds for all group sizes. Figure 4 illustrates these considerations and the following Proposition 4.

**Figure 4:**
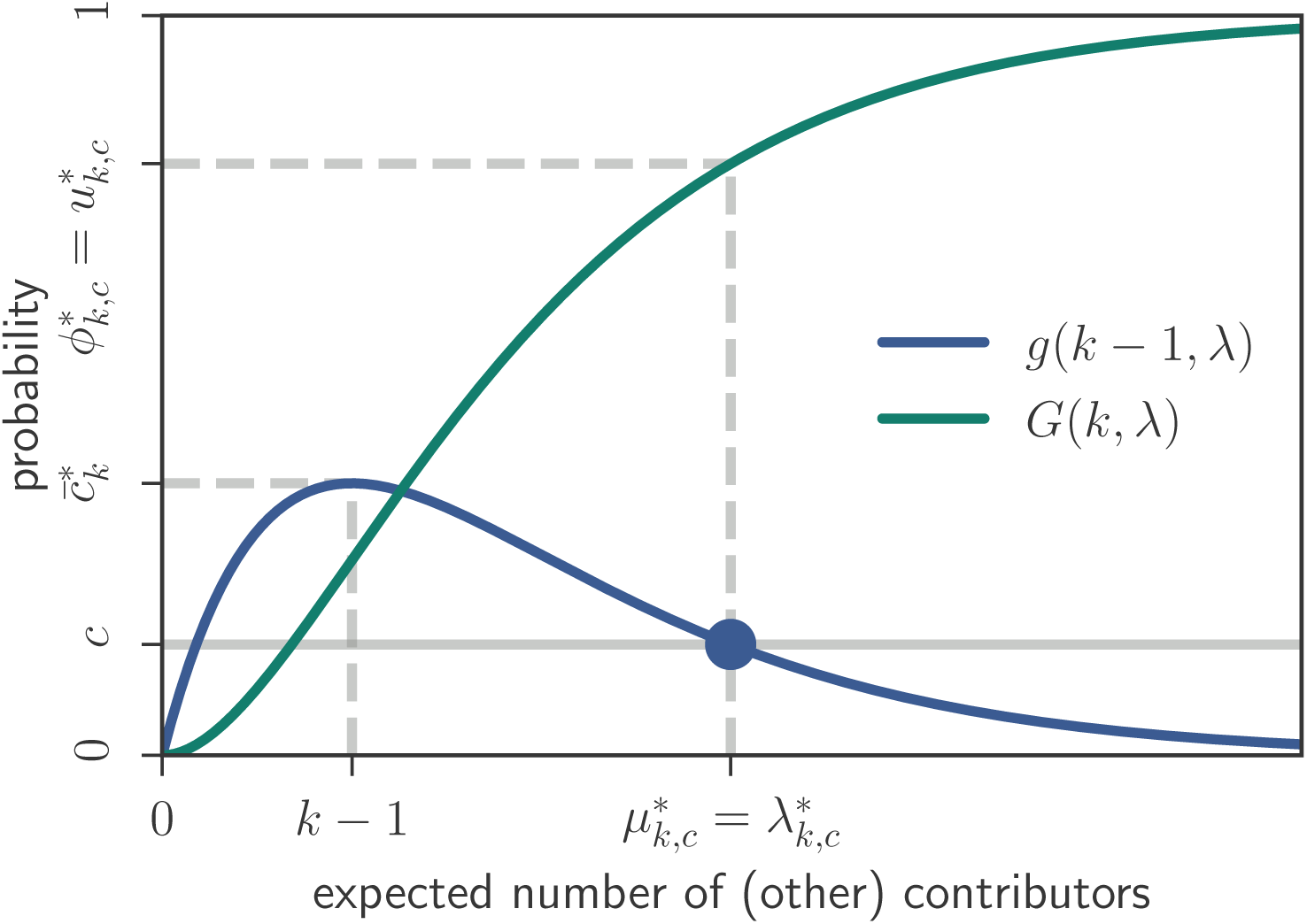
Illustration of Lemma 3 and Proposition 4 for *k* = 2 and *c* = 0.2. The limit of the expected number of (other) contributors 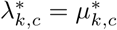 is given by the larger of the two solutions to the pivotality condition *g*(*k* − 1, *λ*) = *c*. Both the equilibrium payoff 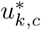 and the success probability 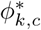 converge to the probability 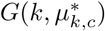 that there are at least *k* contributors given that the number of contributors follows a Poisson distribution with expected value 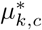.

From the convergence of *λ*_*k,c*_(*n*) = (*n* − 1) · *p*_*k,c*_(*n*) to the finite limit 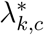 it is immediate that 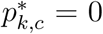 holds. This in turn implies that the expected number of contributors converges to the same limit as *λ*_*k,c*_(*n*), that is, 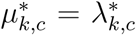 holds. To prove the following result it remains to establish that both the success probability and the equilibrium payoff converge to the probability that there are at least *k* contributors given that the number of contributors follows a Poisson distribution with expected value 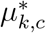. This probability is 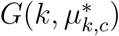, where

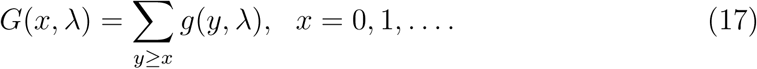

#### Proposition 4.

*Let 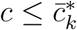. Then*,

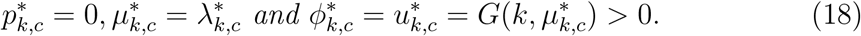

*Proof.* Using a generalization of the classical Poisson approximation (see, for instance, Billingsley, 1995, Theorem 23.2) we have that (i) 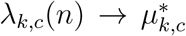 implies 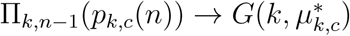 and (ii) 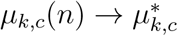 implies 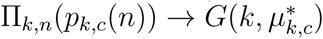. Using (7) and the second equality in (10), this proves the final two equalities in (18). Further, as 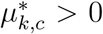 is implied by the equality 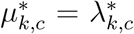 and Lemma 3, we have that, as asserted in (18), the inequality 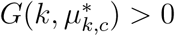 holds. □

We note that the limit values for the expected number of contributors 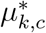 and the success probability 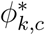 identified in Proposition 4 coincide with the values one would obtain for the maximal equilibrium in a Poisson game (Myerson, 1998) in which group size is commonly known to be a Poisson random variable with mean 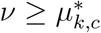 but the actual realization of the group size is unknown to individuals. This holds for any *µ* satisfying the above inequality, that is, in the Poisson game the expected number of contributors and the success probability are independent of expected group size as long as the expected group size is large enough. These observations are in line with the convergence results from Makris (2009), who considers a model for the provision of a binary public good in which, in contrast to the scenario we consider, there is cost-sharing between participants and unused contributions are returned.

It is of interest to ask under which circumstances the expected number of contributors *µ*_*k,c*_(*n*) exceeds the number of contributors *k* that are required for the provision of the public good (Gradstein and Nitzan, 1990, Section 4). Lemma 3 and Proposition 4 provide an answer to this question for large group sizes. As the probability mass function *g*(*k, λ*) of the Poisson distribution is decreasing in its parameter *λ* for *λ* ≥ *k* (see the proof of Lemma 3) the following holds: for small cost (0 < *c* < *g*(*k* − 1, *k*)) the expected number of contributors exceeds *k* for sufficiently large groups, whereas for intermediate cost (*g*(*k* − 1, *k*) < *c* ≤ *g*(*k* − 1, *k* − 1)) the reverse is true.

### 3.5. The effect of group size on the expected number of contributors in large groups

As we have discussed for the equilibrium payoff *u*_*k,c*_(*n*) (and is also the case for the success probability *ϕ*_*k,c*_(*n*)), an increase in group size has two countervailing effects on the expected number of contributors *µ*_*k,c*_(*n*) = *n* · *p*_*k,c*_(*n*): on one hand, for a given participation probability an increase in *n* causes the expected number of contributors to increase; on the other hand, an increase in *n* causes *p*_*k,c*_(*n*) to fall (Proposition 1). In light of Propositions 2 and 3 it is natural to conjecture that the second of these effects dominates (i.e., that the expected number of contributors is strictly decreasing in group size). However, the following proposition shows that this is not necessarily the case. Specifically, Proposition 5 shows that for sufficiently large groups the comparative statics of the expected number of contributors are determined by how the participation cost *c* compares to a critical cost level, given by 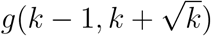: for costs below this level the expected number of contributors is strictly increasing in group size, whereas for costs above this level (but low enough for non-trivial equilibria to exist for all group-sizes) the expected number of contributors is strictly decreasing in group size.

#### Proposition 5.

*(i) Suppose 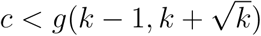 or, equivalently, 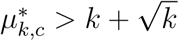 holds. Then there exists N such that µ*_*k,c*_(*n* + 1) > *µ*_*k,c*_(*n*) *holds for all n* > *N.*

*(ii) Suppose 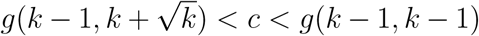 or, equivalently, 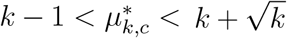 holds. Then there exists N such that µ*_*k,c*_(*n* + 1) < *µ*_*k,c*_ (*n*) *holds for all n* > *N*.

The proof of Proposition 5 is in Appendix A.5. It uses Proposition 4 only to ensure the equivalences noted in the statement of Proposition 5. In particular, the proof does *not* use Proposition 4 to approximate the binomial probabilites appearing in the pivotality condition (2) by their Poisson counterparts. Instead, our proof relies on inequalities for the probability mass function of the binomial distribution established in Anderson and Samuels (1967). This is essential, as the arguments presented in the proof of Proposition 7(iii) in Hindriks and Pancs (2002) show that using the standard Poisson approximation to the pivotality condition for finite *n* leads to the mistaken conclusion that for sufficiently large *n* the expected number of contributors is decreasing in group size irrespectively of the participation cost.

## 4. Discussion

We have studied the comparative statics of the maximal symmetric equilibrium in the class of participation games without refunds introduced in Palfrey and Rosenthal (1984). We have found that for all thresholds *k* > 1 the probability of participation, the expected payoff, and the probability that the public good is provided in this equilibrium are strictly decreasing in group size if the participation cost *c* is no larger than the critical value 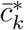 identified in equation (4). Otherwise, these results hold for all group sizes no larger than a critical group size 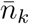 with the participation probability, the equilibrium payoff, and the probability that the public good is provided all being zero for group sizes exceeding 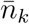. In line with the suggestion from Palfrey and Rosenthal (1984, p. 178) that “mixed strategy equilibria ‘disappear’ as the number of players grows large” we have found that the probability of participation converges to zero as group size goes to infinity. However, for cost 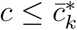 this does not imply that the equilibrium payoff and the probability of provision converge to zero. Rather, both of these converge to the same strictly positive limit that has a simple characterization in terms of the Poisson distribution. We have also signed the group size effect on the expected number of contributors for large groups, showing, in particular, that this effect is positive for sufficiently small cost. Overall, these results provide an almost complete picture of the effects of group size on the maximal symmetric equilibria in the class of games that we have studied.^10^

In their pioneering work, Palfrey and Rosenthal (1984) also consider a second version of their participation game in which contributions are refunded when the number of contributors fails to reach the necessary threshold. As shown in Palfrey and Rosenthal (1984) this participation game with refunds has a unique non-trivial symmetric equilibrium *q*_*k,c*_(*n*). For all *c* ∈ (0, 1) and *n* > *k* this equilibrium satisfies 0 < *q*_*k,c*_(*n*) < 1 and the indifference condition

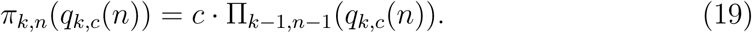

Despite the significant structural difference between the participation games without and with refunds, our arguments can be adapted to show that for 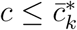 counterparts to Propositions 1 – 3 hold for the participation game with refunds. We show this in Appendix A.6.^11^

Makris (2009) considers symmetric equilibria in a variant of the participation games with refunds in which excessive contributions are also refunded when the threshold is passed: when there are 𝓁 > *k* participants the public good is provided, but each participant only pays the cost *ck*/𝓁. While his main focus is on games in which there is both uncertainty about preferences and group size, he also briefly considers (in his Section 3) the counterpart to the symmetric complete information model that we have studied here and suggests in passing “that for sufficiently large group-size the expected number of contributors and the probability of provision decreases with group-size” (Makris, 2009, p. 296) in the unique non-trivial symmetric equilibrium of such a model. In our view, it would be interesting to verify these claims and check whether arguments similar to the ones we have given here can be used to extend them to all group sizes.

Much of the recent literature on participation games has focused on the (more realistic) case, first considered in Palfrey and Rosenthal (1988), in which there is incomplete information about costs and/or benefits. The comparative statics in such models are quite different from the ones in the complete information model that we have considered here. For instance, Johnson (2002) considers group-size effects in a version of the volunteer’s dilemma featuring private information both about idiosyncratic costs and benefits and finds that — contrary to what is true in the volunteer’s dilemma with complete information and our model — the effect of group size on the success probability is ambiguous whereas equilibrium payoffs are strictly increasing in group size. Palfrey and Rosenthal (1988) characterize (see their Table 2) the comparative statics of the participation probability with respect to group size when agents have private information about their idiosyncratic altruistic “warm glow” benefit from contributing to the public good, with Goeree and Holt (2005, Proposition 4 and Footnote 20) then obtaining conditions under which this characterisation implies that the participation probability first falls and then rises with an increase in group size. In light of such results, there is little hope of generalizing our arguments to participation games with incomplete information.

## Appendix

### A.1. Proof of Lemma 2

Using the definition in (1), straightforward algebra shows that the equation *π*_*k,n*_(*p*) = *π*_*k,n*+1_(*p*) has a unique solution in the interval (0, 1) given by 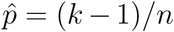. From the unimodality properties of the pivot probability *π*_*k,n*_(*p*) noted in Section 2.2, (*k* − 1)/(*n* − 1) is the unique maximizer of *π*_*k,n*_(*p*) over *p* ∈ (0, 1), so that *π*_*k,n*_((*k* − 1)/(*n* − 1)) > *π*_*k,n*_((*k* − 1)*/n*) holds. Thus, we have *π*_*k,n*_((*k* − 1)/(*n* − 1)) > *π*_*k,n*+1_((*k* − 1)*/n*). Recalling the definition of 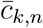 in (3) we thus have 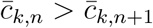.

From equations (1) and (3) we have, upon setting *m* = *n* − 1,

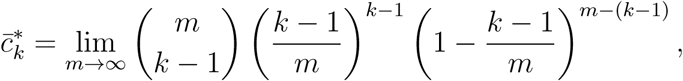

so that (4) follows from the classical Poisson approximation to the binomial distribution.

### A.2. Proof of Proposition 1

Let *P* := ((*k* − 1)/(*n* − 1), 1) and *Q* := [(*k* − 1)*/n*, 1). As *π*_*k,n*_((*k* − 1)/(*n* − 1)) > *π*_*k,n*+1_((*k* − 1)*/n*) holds (Lemma 2), it follows from the unimodality properties of the pivot probabilities that for all *q* ∈ *Q* there exists a unique *h*(*q*) ∈ *P* such that

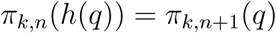

holds. Further, the function *h* : *Q* → *P* thus defined is continuous (in fact, differentiable, as *π*_*k,n*+1_(*q*) is differentiable on *Q* and the inverse of the restriction of *π*_*k,n*_(*p*) to the interval *P* is differentiable by the inverse function theorem). Observing that *h*((*k* − 1)*/n*) > (*k* − 1)/(*n* − 1) > (*k* − 1)*/n* holds, where the first inequality is from *h*((*k* − 1)*/n*) ∈ *P*, and that *π*_*k,n*_(*p*) and *π*_*k,n*+1_(*p*) have no intersection in the interval *P* (see the proof of Lemma 2), we obtain *h*(*q*) > *q* for all *q* ∈ *Q*.

The condition 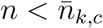 in the statement of the proposition implies 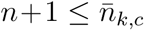 and therefore that non-trivial equilibria exist for group sizes *n* and *n* + 1. Thus, Lemma 1 yields *p*_*k,c*_(*n*) ∈ *P, p*_*k,c*_(*n* + 1) ∈ *Q*, and *π*_*k,n*_(*p*_*k,c*_(*n*)) = *π*_*k,n*+1_(*p*_*k,c*_(*n* + 1)) = *c* > 0, so that *p*_*k,c*_(*n*) = *h*(*p*_*k,c*_(*n* + 1)) holds. Consequently, *p*_*k,c*_(*n*) > *p*_*k,c*_(*n* + 1) follows from the inequality *h*(*q*) > *q* established in the preceding paragraph.

### A.3. Proof of Proposition 2

As in the proof of Proposition 1, let *P* = ((*k* − 1)/(*n* − 1), 1), *Q* = [(*k* − 1)*/n*, 1), and let *h* : *Q* → *P* denote the continuous function satisfying

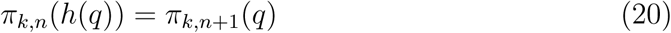

for all *q* ∈ *Q*. Following reasoning analogous to the one in the proof of Proposition 1 and using (10) to rewrite the equilibrium payoffs it suffices to show

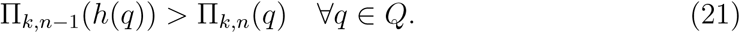

Towards this end, let us define

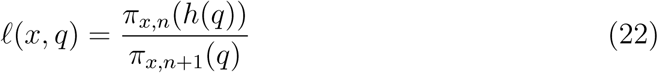

for *q* ∈ *Q* and *x* = 1, …, *n*. From (20) we have *𝓁*(*k, q*) = 1 for all *q* ∈ *Q*. In the following we first argue that this implies *𝓁*(*k* + 1, *q*) > 1 and then, in a second step, show that this inequality implies (21). The key observation underlying these arguments is due to Klenke and Mattner (2010), who (in the proof of their Lemma 2.4) observe that, for *x* = 1, …, *n* − 1,

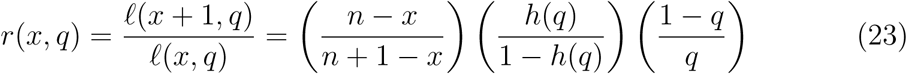

is decreasing in *x*. The second equality in (23) follows from (1) and (22).

Step 1: We show *𝓁*(*k* + 1, *q*) > 1 for all *q* ∈ *Q*. Because *𝓁*(*k, q*) = 1, this is equivalent to showing that *r*(*k, q*) > 1 holds for all *q* ∈ *Q*.

It will be useful to begin by establishing the inequality *r*(*k, q*) > 1 for the lower endpoint of the interval *Q*, that is, *q* = (*k* − 1)*/n*. Substituting this value for *q* into equation (23) yields

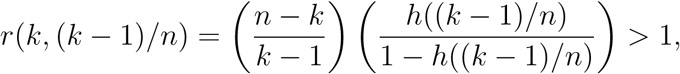

where the inequality is implied by *h*((*k* − 1)*/n*) > (*k* − 1)/(*n* − 1). Now suppose there exists *q* ∈ ((*k* − 1)*/n*, 1) satisfying *r*(*k, q*) ≤ 1. Because *h* : *Q* → *P* is continuous in *q*, so is *r*(*k, q*). By the intermediate value theorem for continuous functions there then exists 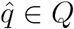 satisfying 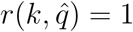. As 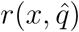 is decreasing in *x*, this implies 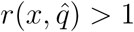 for all *x* satisfying 1 ≤ *x* < *k*. Consequently, 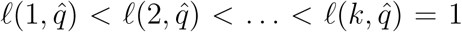 holds, where the equality is from (20) and (22). Similarly, we have 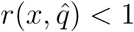 for all *x* satisfying *k* < *x* ≤ *n* − 1, which implies 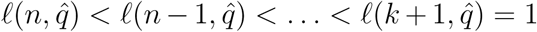, where the equality is from 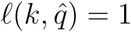 and 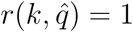. Hence, we have 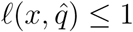 for *x* = 1, …, *n* with strict inequality for *x* ∉ {*k, k* + 1}. Consequently, from (22) (and the assumption *k* > 1) we have

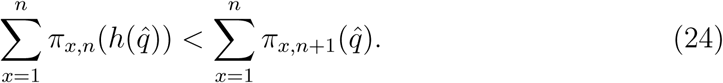

But this is impossible: from (1) the left side of (24) is one, whereas the right side is smaller than one. Hence, no 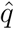 satisfying 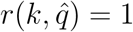 exists and we have *𝓁*(*k* + 1, *q*) > 1 for all *q* ∈ *Q*.

Step 2: From *𝓁*(*k* + 1, *q*) > 1 and *𝓁*(*k, q*) = 1 we have *r*(*k, q*) > 1. As *r*(*x, q*) is decreasing in *x*, this implies *r*(*x, q*) > 1 for all *x* satisfying 1 ≤ *x* ≤ *k*. Consequently, *𝓁*(1, *q*) < *𝓁*(2, *q*) < … < *𝓁*(*k, q*) = 1 holds. Because we have assumed *k* > 1 this implies

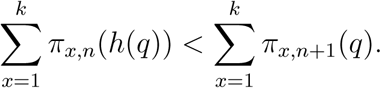

From (1) and (7) this is equivalent to (21).

### A.4. Proof of Lemma 3

Upon taking logarithms in (5) it is easily verified that *g*(*k* − 1, *λ*) is differentiable and decreasing in *λ* on [*k* − 1, ∞) with lim_*λ*→∞_ *g*(*k* − 1, *λ*) = 0. Hence, as asserted in the statement of the lemma, the condition *g*(*k* − 1, *λ*) = *c* has a unique solution satisfying *λ* ≥ *k* − 1 that we denote by 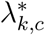. It remains to show that *λ*_*k,c*_(*n*) converges to this value as *n* → ∞.

As *p*_*k,c*_(*n*) satisfies the pivotality condition (2) for all *n*, we have

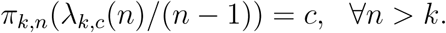

From Lemma 1 we also have the inequality *λ*_*k,c*_(*n*) > *k* − 1 for all *n* > *k*.

Let ϵ > 0 and 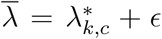. Then 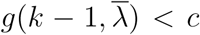 holds and, by the Poisson approximation to the binomial distribution, there exists *N*_1_ such that 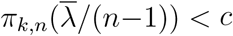 holds for all *n* > *N*_1_. From the unimodality properties of the pivot probability *π*_*k,n*_(*p*), we then have that 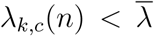 holds for all *n* > *N*_1_. Let 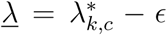. If *λ* > *k* − 1 holds, then, using an analogous argument to the one we used when considering 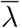, there exists *N*_2_ such that *λ*_*k,c*_(*n*) > *λ* holds for all *n* > *N*_2_. If *λ* ≤ *k* − 1 holds, then define *N*_2_ = *k*. We then again have that *λ*_*k,c*_(*n*) > *λ* holds for all *n* > *N*_2_. Letting *N* = max{*N*_1_, *N*_2_} we have established that for all *e* > 0 there exists *N* such that for all *n* > *N* the inequalities 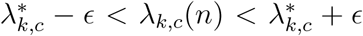 are satisfied. Consequently, *λ*_*k,c*_(*n*) converges to 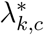.

### A.5. Proof of Proposition 5

*Proof.* (i) Suppose 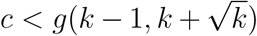 holds. From Proposition 4 and Lemma 3 we have that 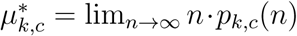 satisfies 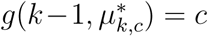 and 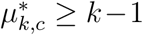. Because *g*(*k* − 1, *λ*) is decreasing in *λ* for *λ* ≥ *k* − 1 (see the beginning of the proof of Lemma this implies 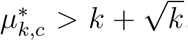. An analogous argument shows that 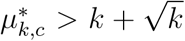 implies 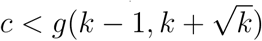.

Suppose 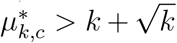 holds. As *µ*_*k,c*_(*n*) = *n* · *p*_*k,c*_(*n*) converges to 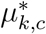 there thus exists *N* such that 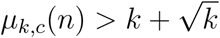 holds for all *n* > *N*. Consider any such *n*. The first part of Theorem 3.1 in Anderson and Samuels (1967) then implies

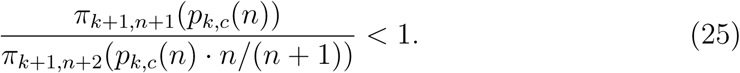

Simple algebra shows

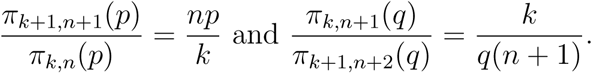

Thus,

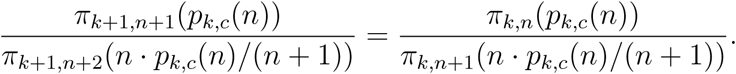

Hence, (25) implies

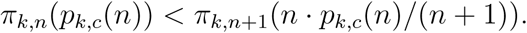

Because *π*_*k,n*_(*p*_*k,c*_(*n*)) = *π*_*k,n*+1_(*p*_*k,c*_(*n* + 1)) = *c* holds (as both *p*_*k,c*_(*n*) and *p*_*k,c*_(*n* + 1) are non-trivial), we thus have

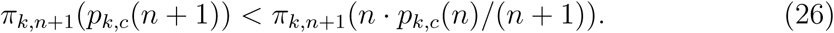

To establish the first part of the proposition, it remains to show that this inequality implies

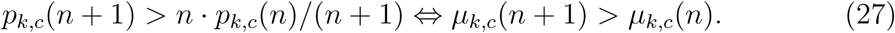

The pivot probability *π*_*k,n*_(*p*) is strictly decreasing in *p* in [(*k* − 1)*/n*, 1] and Lemma 1 implies that *p*_*k,c*_(*n* + 1) ≥ (*k* − 1)*/n* holds. Thus, a violation of the first inequality in (27) contradicts the inequality in (26), so that the desired conclusion follows.

(ii) The equivalence 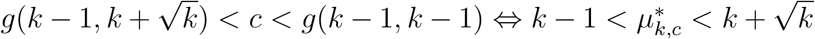 follows as the equivalence in part (i).

Suppose 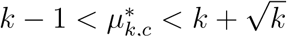 holds. Fix *λ* and 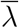 such that

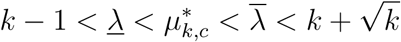

holds. Because *µ*_*k,c*_(*n*) = *n* · *p*_*k,c*_(*n*) converges to 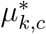 there exists 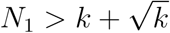 such that 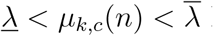 holds for all *n* > *N*_1_. Consider any such *n*. The function

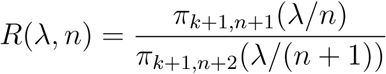

is unimodal in *λ* in 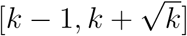.^12^ Thus, the inequality

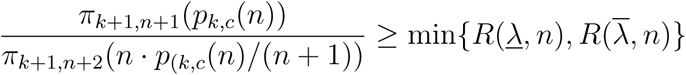

holds. Observing that 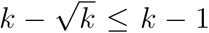 holds, the second part of Theorem 3.1 in Anderson and Samuels (1967) implies that there exists *N*_2_ ≥ *N*_1_ such that for all *n* > *N*_2_ the inequality 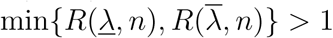 is satisfied. Hence, for all *n* > *N*_2_ we have

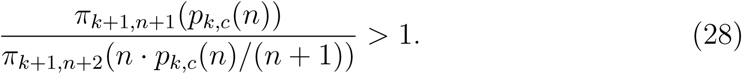

Arguments analogous to the ones showing that (25) implies part (i) of the proposition show that (28) implies *p*_*k,c*_(*n* + 1) < *n* · *p*_*k,c*_(*n*)/(*n* + 1), proving part (ii) of the proposition.

□

### A.6. Participation games with refunds

Suppose 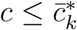, so that 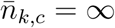 and Propositions 1 – 3 hold for all *n* > *k* > 1. The following arguments show that corresponding results then hold for the unique non-trivial symmetric equilibrium of the participation game with refunds.

We first show that the participation probability is strictly decreasing in group size, that is, that *q*_*k,c*_(*n* + 1) < *q*_*k,c*_(*n*) holds for all *n* > *k*. From Proposition 6 in Palfrey and Rosenthal (1984) we have *q*_*k,c*_(*n*) > *p*_*k,c*_(*n*) for all *n*. Thus, using the same notation as in the proof of Proposition 1 for the sets *P* and *Q*, we have *q*_*k,c*_(*n*) ∈ *P* and *q*_*k,c*_(*n* + 1) ∈ *Q* (because the equilibria *p*_*k,c*_(*n*) and *p*_*k,c*_(*n* + 1) are both non-trivial by the assumption 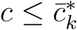). Now suppose that *q*_*k,c*_(*n* + 1) ≥ *q*_*k,c*_(*n*) holds. As *π*_*k,n*+1_(*p*) < *π*_*k,n*_(*p*) holds on *P* and *π*_*k,n*+1_(*p*) is strictly decreasing on this domain, *q*_*k,c*_(*n* + 1) ≥ *q*_*k,c*_(*n*) ∈ *P* implies *π*_*k,n*+1_(*q*_*k,c*_(*n* + 1)) < *π*_*k,n*_(*q*_*k,c*_(*n*)). We also have that *q*_*k,c*_(*n* + 1) ≥ *q*_*k,c*_(*n*) > 0 implies Π_*k*−1,*n*_(*q*_*k,c*_(*n* + 1)) > Π_*k*−1,*n*−1_(*q*_*k,c*_(*n*)) as Π_*k*−1,*n*−1_(*p*) is increasing both in *n* and *p*. We thus obtain that the left side of (19) strictly decreases when the group size is increased from *n* to *n* + 1 whereas the right side of (19) strictly increases, contradicting the hypothesis that *q*_*k,c*_(*n* + 1) is the symmetric equilibrium for group size *n* + 1. Hence, *q*_*k,c*_(*n* + 1) < *q*_*k,c*_(*n*) must hold.

Second, we show that the equilibrium payoff is strictly decreasing in group size. Denoting the equilibrium payoff in the participation game without refunds as a function of group size by *v*_*k,c*_(*n*) we have (by the indifference condition) *v*_*k,c*_(*n*) = Π_*k,n*−1_(*q*_*k,c*_(*n*)) for all *n* > *k*. Hence, our task is to show that Π_*k,n*_(*q*_*k,c*_(*n* + 1)) < Π_*k,n*−1_(*q*_*k,c*_(*n*)) holds for all *n* > *k*. Towards this end, observe that (19) can be rewritten as

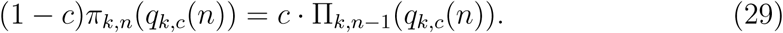

Now suppose that Π_*k,n*_(*q*_*k,c*_(*n* + 1)) ≥ Π_*k,n*−1_(*q*_*k,c*_(*n*)) holds. From (29) we must then have *π*_*k,n*+1_(*q*_*k,c*_(*n* + 1)) ≥ *π*_*k,n*_(*q*_*k,c*_(*n*). As *π*_*k,n*+1_(*q*) is continuous and decreasing with limit *π*_*k,n*+1_(1) = 0 on *Q*, there then exists 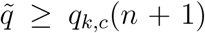 that satisfies 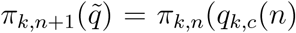. Because *q*_*k,c*_(*n*) ∈ *P* and 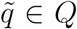 hold, the same argument as in the proof of Proposition 2 implies 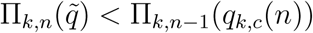. But as Π_*k,n*_(*q*) is increasing in *q*, this contradicts the hypothesis Π_*k,n*_(*q*_*k,c*_(*n* + 1)) ≥ Π_*k,n*−1_(*q*_*k,c*_(*n*)). Hence, Π_*k,n*_(*q*_*k,c*_(*n* +1)) < Π_*k,n*−1_(*q*_*k,c*_(*n*)) must hold, proving that *v*_*k,c*_(*n* +1) < *v*_*k,c*_(*n*) holds.

Third, we conclude the argument by establishing that the success probability is strictly decreasing in *n*, too.

As we are considering a non-trivial equilibrium with *q*_*k,c*_(*n*) > 0, choosing to contribute with probability one is a best response if all other agents contribute with probability *q*_*k,c*_(*n*). A player choosing this strategy obtains the public good and pays the contribution cost if and only if at least *k* − 1 out of the other *n* − 1 group members contribute. Hence, the equilibrium payoff satisfies

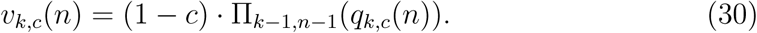

The equilibrium payoff is also given by the probability that the public good is provided if all *n* players contribute with probability *q*_*k,c*_(*n*) minus the expected cost of contribution when following this strategy. As each player contributes with probability *q*_*k,c*_(*n*) and in this case has to pay the cost *c* if and only if at least *k* − 1 of the remaining *n* − 1 players contribute, this means that the equilibrium payoff can also be written as

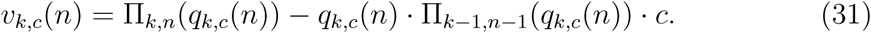

Substituting from (30) into (31) to eliminate Π_*k*−1,*n*−1_(*q*_*k,c*_(*n*)) from the latter equation, we obtain

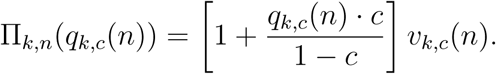

Because both *q*_*k,c*_(*n*) and *v*_*k,c*_(*n*) are strictly decreasing in *n*, it follows that the success probability Π_*k,n*_(*q*_*k,c*_(*n*)) is strictly decreasing in *n*.

## Acknowledgement

We thank Marco Archetti, Christian Kleiber, Marc-Andreas Muendler, Larry Samuelson, and Urs Schweizer for helpful discussions and comments. J. Peña acknowledges IAST funding from the French National Research Agency (ANR) under the Investments for the Future (Investissements d’Avenir) program, grant ANR-17-EURE-0010. The Python code used for creating the figures of this paper is publicly available on GitHub (https://github.com/jorgeapenas/olsonconjecture).

## Footnotes

1 See Sandler (2015) for a summary of Olson’s hypotheses on the effects of group composition and some of the literature investigating such effects.

2 The impossibility of role identification is the standard justification for the focus on the symmetric strategy profiles in symmetric games in evolutionary game theory (see Weibull, 1995). The robustness of such symmetric equilibria against perturbations (which maintain the structure of a threshold public good game but abandon symmetry) is discussed in Kalandrakis (2009), who obtains a purification result along the lines of Harsanyi (1973). See Binmore and Samuelson (2001) for a discussion of the relationship between the purification of mixed strategy equilibria and their evolutionary stability in an environment with noisy role identification. For a more heuristic justification for the focus on the symmetric equilibria in participation games see Dixit and Olson (2000).

3 This is immediate from the fact that, as we note in Section 2.2 and illustrate in Figure 1, the pivot probability, defined in equation (1), is strictly decreasing (increasing) in the participation probability at the larger (smaller) of the two non-trivial symmetric equilibria. See McBride (2006) for informal discussion and Peña et al. (2014), who provide an evolutionary model and characterize the stability properties of symmetric equilibria for a broad class of symmetric binary-action games, for a proof.

4 The main focus of Hindriks and Pancs (2002) is on the effect of altruism on the equilibrium provision of the public good. For the the teamwork dilemma (*k >* 1) we study here, the version of altruism they consider does not affect the probability that the public good is provided and can therefore be ignored (Hindriks and Pancs, 2002, Proposition 4).

5 In the case *k* = *n*, which corresponds to an assurance problem in the sense of Sen (1967), it is an equilibrium for all group members to participate, thereby ensuring the provision of the public good. We exclude this case as it adds nothing of interest to our analysis in Section 3 but would require an additional case distinction.

6 The game has a multitude of asymmetric Nash equilibria, including those in which the players coordinate in such a way that *k* players contribute and the remaining *n* − *k* players do not contribute. Further asymmetric equilibria are described in Palfrey and Rosenthal (1984, Section 2).

7 The first two of these properties are immediate from the definition of *π*_*k,n*_(*p*) in (2). The remaining properties follow from observing that 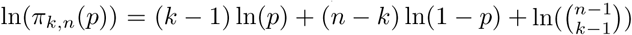 is strictly concave in *p* on (0, 1) and has its unique critical point at *p* = (*k* 1)/(*n* − 1).

8 Indeed, as we have shown in previous work (viz., Peña and Nöldeke, 2018), for a large class of symmetric participation games it is the case that *all* interior symmetric equilibria feature participation probabilities that are strictly decreasing in group size.

9 Otherwise, the critical group size 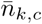 is finite and it is immediate from (13) that all of the limits defined above are equal to zero.

10 The one missing piece is a result characterizing the group size effect on the expected number of contributors for all (rather than only for large) group sizes. We conjecture that there are only three possibilities, namely that the expected number of contributors is (i) decreasing throughout, (ii) increasing throughout, or (iii) unimodal. Proving this conjecture is a non-trivial task as it requires to extend the analysis from Anderson and Samuels (1967) to obtain a complete characterization of the comparative statics of the binomial probability mass function when the sample size and the success probability are changed in such a way that the expected number of successes stays constant.

11 The role of the condition 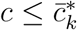 in this proof is to ensure that *q*_*k,c*_(*n*) exceeds the mode (*k* − 1)/(*n* − 1) of the pivot probability *π*_*k,n*_(*p*) for all *n*. Whether counterparts to Propositions 2 and 3 also hold for the model with refunds when the participation cost exceeds *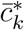* is an open question.

12 To verify this claim calculate 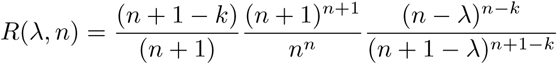 and observe that 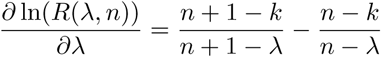 is positive for *λ* ∈ [*k* − 1, *k*) and negative for 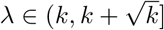.

